# Generative machine learning of ADAR substrates for precise and efficient RNA editing

**DOI:** 10.1101/2024.09.27.613923

**Authors:** Yue Jiang, Lina R. Bagepalli, Bora S. Banjanin, Yiannis A. Savva, Yingxin Cao, Lan Guo, Adrian W. Briggs, Brian Booth, Ronald J. Hause

## Abstract

Adenosine Deaminase Acting on RNA (ADAR) converts adenosine to inosine within certain double-stranded RNA structures. However, ADAR’s promiscuous editing and poorly understood specificity hinder therapeutic applications. We present an integrated approach combining high-throughput screening (HTS) with generative deep learning to rapidly engineer efficient and specific guide RNAs (gRNAs) to direct ADAR’s activity to any target. Our HTS quantified ADAR-mediated editing across millions of unique gRNA sequences and structures, identifying key determinants of editing outcomes. We leveraged these data to develop DeepREAD (Deep learning for <u>R</u>NA <u>E</u>diting by <u>A</u>DAR <u>D</u>esign), a diffusion-based model that elucidates complex design rules to generate novel gRNAs outperforming existing design heuristics. DeepREAD’s gRNAs achieve highly efficient and specific editing, including challenging multi-site edits. We demonstrate DeepREAD’s therapeutic potential by designing gRNAs targeting the MECP2^R168X^ mutation associated with Rett syndrome, achieving both allelic specificity and species cross-reactivity. This approach significantly accelerates the development of ADAR-based RNA therapeutics for diverse genetic diseases.

---

Adenosine deaminase acting on RNA (ADAR) is a ubiquitously expressed enzyme that converts adenosine to inosine within double-stranded RNA (dsRNA) ^1^. Inosine mimics guanosine during splicing and translation, allowing ADAR to be used therapeutically to modulate adenosine-containing regulatory RNA sequences and recode amino acids ^2^. ADAR’s ability to recode transcripts in vertebrates to alter protein function has been well-documented, with notable examples in neurotransmitter receptors and innate immunity regulation ^3, 4^. For instance, ADAR edits the family of GRIA receptors in the nervous system, altering calcium permeability ^5^. Additionally, ADAR’s promiscuous deamination of adenosines within dsRNA structures marks transcripts as “self”, preventing stimulation of the innate immune system upon cytoplasmic export ^4, 6^.

Humans possess two catalytically active ADAR isoforms: ADAR1 and ADAR2. ADAR1 can produce two variants determined by their transcriptional start site: the ubiquitously expressed ADAR1 p110 and the interferon-induced p150 ^7^. ADAR1 p110 and ADAR2 localize to the nucleus, while ADAR p150 is cytoplasmic. ADAR2 expression is primarily restricted to the brain and heart ^8^. Both isoforms are essential for normal development and function, as demonstrated by knockout studies in mice ^9^. ADAR exhibits known sequence preferences, such as a 5′-UAG-3′ motif ^10, 11^; however, the general principles governing its specificity remain poorly understood. Recent structural studies have provided insights into ADAR’s interaction with RNA substrates, revealing a footprint of at least 25 base pairs downstream and 5 base pairs upstream of the target adenosine ^12, 13^. Natural ADAR substrates often contain mismatches, wobble base pairs, bulges, and loops interspersed throughout their structure^14^. Secondary structures downstream of the edited adenosine may affect the engagement or specificity of the double stranded RNA binding domains (dsRBDs), while structures within a few base pairs of the target adenosine may affect the activity of the deaminase domain ^15, 16^. The 5′ G dispreference of ADAR is the result of a steric clash ^17^, and many natural 5′ G substrates have mismatches, bulges, or loops on the 5′ side of the edited adenosine ^18^.

These insights into ADAR structure and function have informed the design of antisense oligonucleotides (ASO) and guide RNAs (gRNAs) to direct ADAR activity to specific targets ^15, 19^. Existing methods for therapeutic gRNA design rely on basic heuristics, such as incorporating A-G mismatches, U deletions, discontinuous hybridization arms, repetitive bulges, or chemical modifications to reduce bystander editing at neighboring adenosines ^20-23^. However, these strategies often result in reduced on-target editing efficiency ^20^. Moreover, the vast theoretical sequence space makes exhaustive experimental testing impractical.

Previous approaches to identify features predictive of ADAR editing have primarily focused on supervised learning of the proximal nucleotide context for specific targets ^24^. Although these studies have provided insights into ADAR’s natural editing preferences, they have not fully addressed how to design gRNAs that can effectively overcome these preferences. High-throughput saturation mutagenesis screening of natural RNA editing substrates has revealed target-specific characteristics predictive of editing outcomes, but these models have struggled to generalize across diverse targets ^24^.

To address these limitations, we have developed an integrated approach combining high-throughput screening (HTS) with advanced machine learning (ML) techniques. This method allows us to rapidly engineer and evaluate diverse gRNA designs, uncovering complex rules that govern ADAR editing efficiency and specificity. Our HTS enables the evaluation of millions of ADAR substrates in a single reaction.

Leveraging this rich data set of ADAR activity across diverse substrates we were able to develop two generative ML frameworks trained on our large-scale RNA editing data: 1) A target-conditioned convolutional neural network (CNN) that uses activation maximization (ActMax) to generate improved gRNAs for specific targets and 2) a target-agnostic model using bit diffusion that transforms gRNA sequences into one-hot encoded images to enable in silico, de novo design of gRNAs for any target of interest. We call this approach DeepREAD (Deep learning for RNA Editing by ADAR Design). These models significantly expand our ability to design efficient and specific gRNAs for both therapeutic and synthetic biology applications.

Our approach not only enhances our understanding of ADAR biology but also provides a powerful tool for the rational design of gRNAs. This advancement has significant implications for the development of RNA editing therapeutics and expands the potential applications of ADAR-mediated editing. To demonstrate the power of this approach, we applied DeepREAD to design gRNAs to restore translation of the pathogenic MECP2^R168X^ nonsense mutation associated with Rett Syndrome. The resulting gRNAs showcase DeepREAD’s capacity to design therapeutically viable candidates with desired cross-reactivity between mouse and human mutant variants, coupled with mutant allele specificity.

## Results

### High-throughput gRNA screening uncovers complex determinants of ADAR editing

We developed a massively parallel biochemical screening platform to address key challenges in directing ADAR-mediated RNA editing. This high-throughput approach enables exploration of the vast RNA secondary structure landscape for therapeutic gRNA design and synthetic biology applications and generation of the largest training dataset to date for RNA editing, enabling the development of powerful generative machine learning models for gRNA design.

As a model target, we chose the LRRK2 G2019S mutation, a G-to-A substitution highly penetrant in Parkinson’s disease ^25^. Initial testing of a canonical gRNA design (100 nucleotides complementary to the target sequence with a C mismatch opposite the target A) revealed substantial bystander editing, including a prominent bystander edit two nucleotides upstream from the target (the -2 position) (Supplementary Fig. 1). To improve gRNA specificity and generate a diverse dataset, we constructed a library of ∼112,000 gRNAs, randomly sampling mutations within a region spanning 8 nucleotides upstream to 21 nucleotides downstream from the target adenosine.

The library design consisted of the LRRK2 G2019S sequence tethered by a small hairpin to diverse gRNA sequences, with invariant 5′ and 3′ sequences for RT-PCR amplification and sequencing. We transcribed the library into RNA, incubated with recombinant human ADAR1 or ADAR2 (rADAR1 or rADAR2), and then reverse transcribed, PCR amplified and sequenced to quantify A-to-I editing at each adenosine within every RNA substrate (Fig. 1a).

**Fig 1.**
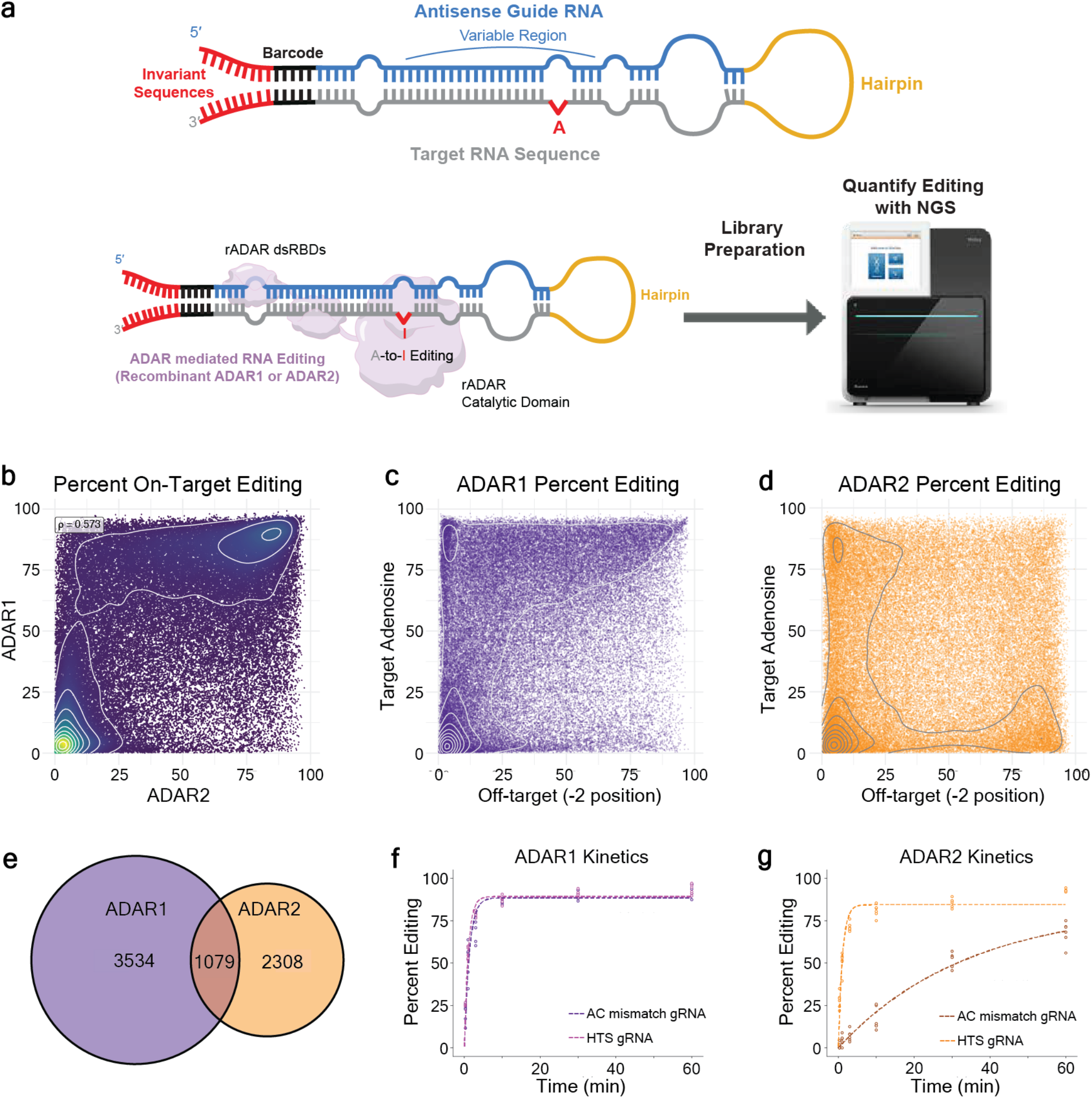
Massively parallel screening of guide RNAs for ADAR-mediated RNA editing. **a**, Schematic of the high-throughput screening platform and LRRK2 G2019S RNA library design. **b**, Correlation between ADAR1 and ADAR2 on-target adenosine editing efficiencies. **c, d**, Correlation between on-target and -2 bystander editing for ADAR1 (**c**) and ADAR2 (**d**). **e**, Identification of highly efficient (on-target editing > 80%) and specific (−2 bystander editing < 20%) gRNA designs for ADAR1, ADAR2, and both isoforms. **f, g**, ADAR editing kinetics measured by time-course experiments in the HTS. Shown are the canonical A-C mismatch and an example substrate with faster kinetics identified from the HTS.

Editing was highly reproducible between technical replicates for both rADAR1 and rADAR2 (ρ > 0.98, Supplementary Fig. 2). However, rADAR1 and rADAR2 on-target editing was less strongly correlated (ρ = 0.57), confirming overlapping but distinct substrate preferences (Fig. 1b) ^10, 26^. We identified numerous gRNAs achieving high on-target editing while minimizing -2 position editing for both ADAR1 and ADAR2 (Fig. 1c, d). Specifically, 4,613 and 3,387 substrates reduced -2 position editing below 20%, while maintaining >80% on-target editing for rADAR1 and rADAR2, respectively, with 1,079 substrates meeting these criteria for both isoforms (Fig. 1e).

To assess deamination kinetics, we conducted time-course experiments on a subset of variants, using the GluR2 R/G hairpin substrate as a control ^27^. GluR2 R/G kinetics matched literature reports ^28, 29^, with rADAR1 slower than rADAR2 (0.16 vs. 1.79 min^-1^) (Supplementary Fig. 3a, b). rADAR2 displayed greater specificity for the R/G target adenosine, while rADAR1 also edited the 5′ neighboring adenosine (Supplementary Fig. 3c). For the LRRK2 canonical design, rADAR1 was unexpectedly faster than for rADAR2 (0.76 vs. 0.03 min^-1^) (Fig. 1f, g). Many HTS designs displayed faster kinetics than the canonical design, indicating that secondary structure influences specificity and deamination rate. One notable design showed similar k_obs_ for ADAR1 and ADAR2 (0.97 and 0.94 min^-1^, respectively) with a predicted secondary structure containing an unusual large internal loop spanning the target adenosine (Fig. 1f, g, h).

These results demonstrate the power of our massively parallel biochemical screen to identify novel dsRNA substrates with improved efficiency and specificity of ADAR-mediated RNA editing, particularly in reducing bystander editing.

### Machine learning models enhance ADAR substrate design

Despite the success of our HTS to identify ADAR substrates with efficient and specific RNA editing of LRRK2 G2019S, current limitations in DNA synthesis and sequencing constrain the fraction of sequence space that can be screened experimentally. We hypothesized that ML could infer the principles governing ADAR editing and explore the expansive sequence space of gRNA design more efficiently than experimental methods alone.

We first developed ML models trained on primary sequence, rADAR1, and rADAR2 editing efficiency, specificity score ((fraction on-target reads + 1) / (sum of bystander reads + 1)), and hairpin melting temperature (T_M_) from our LRRK2 G2019S HTS. Two model architectures were employed: a gradient boosting framework (XGBoost) ^30^ and a CNN ensemble. Both models achieved similar predictive power for editing efficiency, specificity score, and T_M_ (Fig. 2a). While XGBoost offers rapid model building, we focused on the CNN for its capacity to perform generative design of ADAR substrates through activation maximization (ActMax) ^31^.

**Fig 2.**
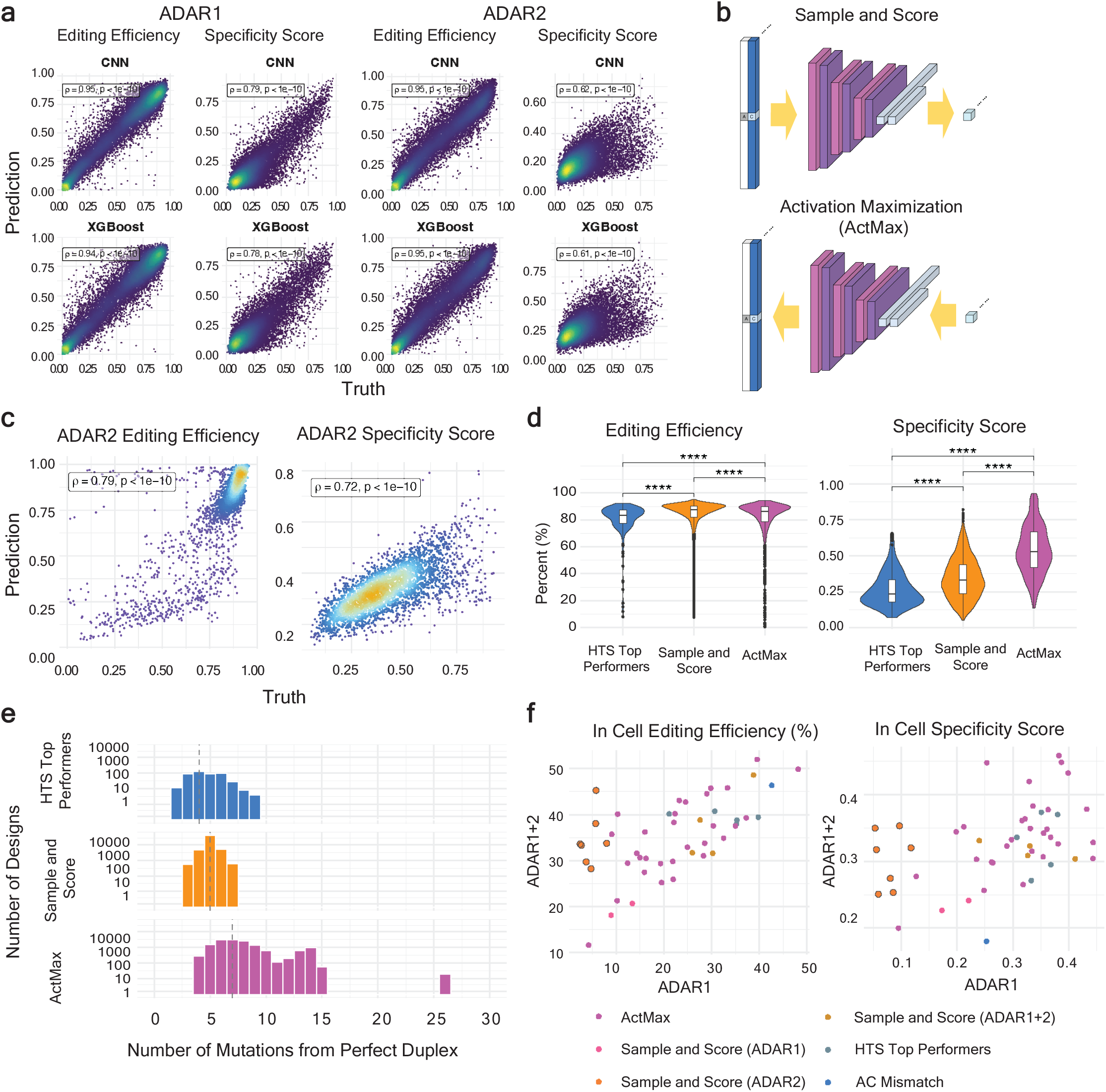
Machine learning models enhance ADAR substrate design. **a**, Predictive performance of CNN ensembles and XGBoost models on held-out test data for editing efficiency and specificity score for ADAR1 and ADAR2, using the LRRK2 HTS data. **b**, Schematic for CNN ensemble-based approaches for generating novel gRNA designs: sample-and-score and activation maximization (ActMax). **c**, Correlation between predicted and observed editing outcomes for LRRK2 generative gRNA designs. **d**, Comparison of editing outcomes between top performers from the original HTS, sample and score designs, and ActMax designs. **e**, gRNA sequence space explored by each approach, represented by the number of mutations from a perfect duplex. **f**, In-cell editing outcomes of selected gRNAs designed via different strategies. Sample and score identified gRNAs specific for the ADAR2 isoform (yellow with grey outline). In **a** and **c**, Spearman’s rho is shown. For **d**, Wilcoxon rank-sum test was applied.

To validate our ML approaches, we generated a new library of ADAR substrates for the LRRK2 G2019S target adenosine using two strategies: ActMax and sample-and-score. ActMax optimized novel gRNA sequences by inputting desired editing metrics into the CNN and iteratively improving the sequences through backpropagation. The sample-and-score method exhaustively mutated gRNA sequences up to a 7-nucleotide Hamming distance from the A-C mismatch in silico and scored variants using the CNN model (Fig. 2b). We included 768 ActMax designs and 2,513 sample-and-score designs in the new library for empirical validation via the HTS.

The ML models demonstrated strong predictive power, yielding Spearman’s r of 0.80 and 0.72 for on-target editing and specificity score, respectively (Fig. 2c). Notably, designs from both approaches outperformed the top substrates from the initial HTS in editing efficiency and specificity, with ActMax-derived gRNA designs achieving the highest specificity (Fig. 2d). ActMax designs showed greater sequence diversity compared to the initial library and sample-and-score designs (Fig. 2e), indicating this method explored a novel sequence space outside original experimental constraints.

To assess the performance of select dsRNA substrates from the HTS as gRNAs in cells, we selected high-performing designs and extended their length to 113 nucleotides for robust endogenous ADAR recruitment and editing ^32, 33^ and were expressed under the control of a novel snRNA expression scaffold ^34^. To mitigate potential increases in bystander editing associated with utilizing longer gRNAs, we insulated the HTS validated sequences between two large symmetrical loops, a strategy previously shown to restrict ADAR activity proximal to the target adenosine, within dsRNA substrates ^23, 35^ (Supplementary Fig. 4). Most of the HTS and ML-derived gRNAs demonstrated significantly better editing specificity than the canonical A-C mismatch in cells (Fig. 2f). Additionally, ActMax generated two gRNAs that outperformed all other methods for enhanced on-target editing efficiency (Fig. 2f).

Our model also identified gRNAs that exhibit ADAR isoform-specific editing. A subset of gRNA designs predicted to drive editing by ADAR2, but not ADAR1, showed enhanced editing in HEK293 cells engineered to express ADAR2, with little to no editing in wild-type (WT) HEK293 cells expressing only endogenous ADAR1. Notably, these gRNAs contain a 5′ G opposite the target adenosine, a feature traditionally considered unfavorable for ADAR editing (Supplementary Fig. 1c). This observation underscores the ability of our models to distinguish between ADAR isoforms and identify novel gRNAs that challenge conventional design principles.

Our findings highlight the potential to overcome the limitations of current heuristics for ADAR-mediated gRNA design, which currently rely on simple rules such as the use of an A-C mismatch or G-G mismatches in a 5′ G context to drive efficient editing of a target adenosine, and/or large internal loops, U-deletions and wobbles to reduce bystander editing ^15, 16, 23^. The success of our ML approaches suggests that a deeper understanding of the principles that govern ADAR editing require large, diverse datasets and sophisticated models.

### Multi-site RNA editing achieved through machine learning

Building on our success with the LRRK2 G2019S target, we next tackled a more challenging target sequence to test the limits of our HTS and ML approach. We sought to leverage ADAR’s promiscuity to selectively edit multiple adenosines within a given dsRNA substrate ^36^ to expand the therapeutic potential of ADAR-mediated RNA editing. We chose to edit a sequence encodingthe BACE1 cleavage site within the transcript encoding the Amyloid Beta Precursor Protein (APP). APP variants that reduce BACE1 have been reported to be protective against Alzheimer’s disease ^37^. Within the coding sequence of the cleavage site, there is a stretch of four nucleotides containing three adenosines, two of which fall within a 5′ G context. Linked RNA editing of these three sites within the same RNA molecule would lead to two amino acid substitutions: K670G and M671V(Fig. 3a).

**Fig 3.**
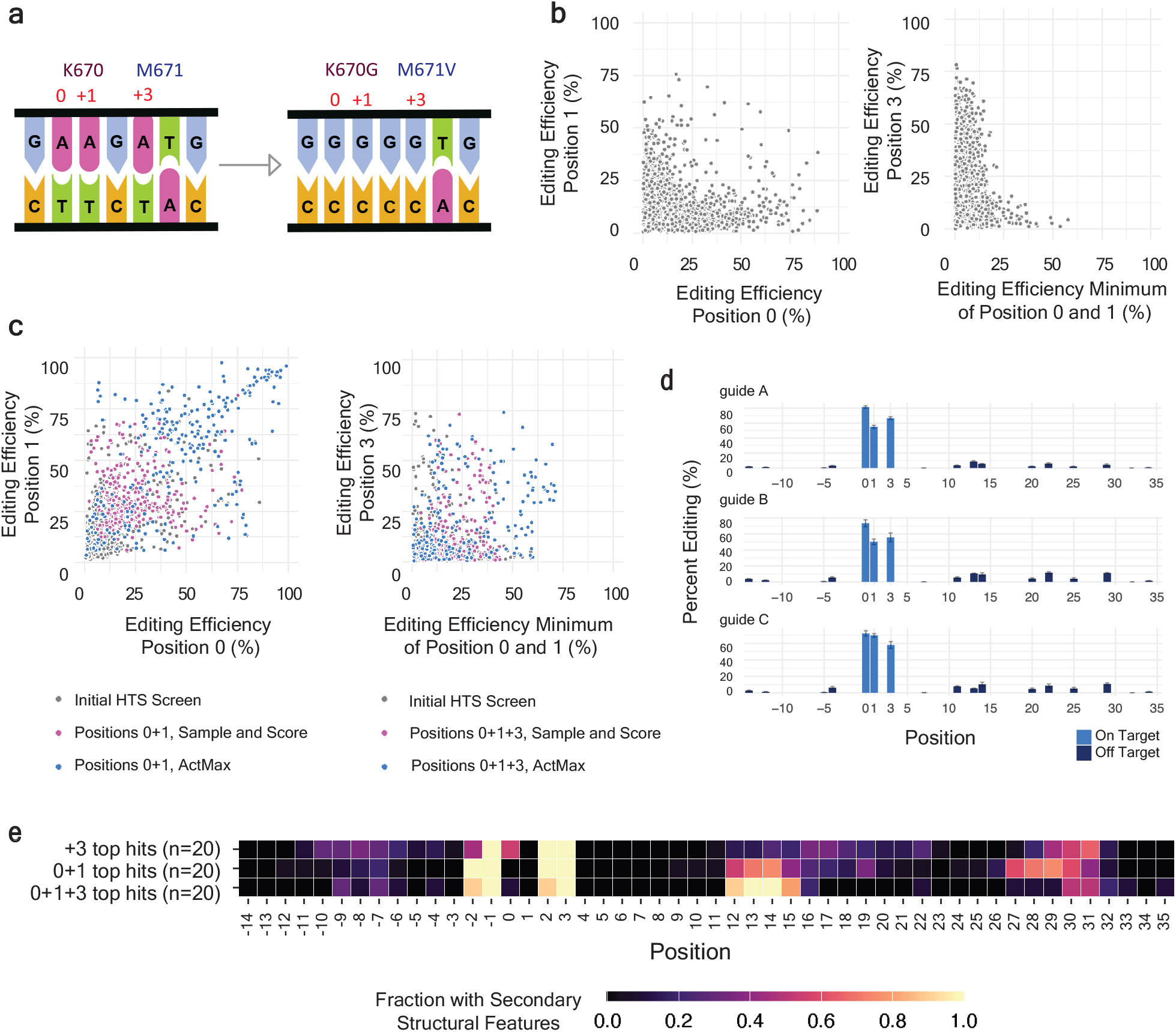
Multi-site editing of ADAR substrates. **a**, Schematic of desired functional alteration of APP, involving two amino acid substitutions (K670G and M671V) that require co-editing of three adenosines, two of which fall within a 5′ G context. **b**, HTS results showing gRNAs with efficient co-editing of 0 and +1 positions (K670G), but no solutions for efficient co-editing of all three sites. **c**, Generative gRNA designs, particularly those from ActMax (blue), improved co-editing of 0 and +1 positions and identified solutions for efficient editing of all three adenosines. **d**, Representative editing profiles of three ActMax-generated gRNA designs showing efficient co-editing of all three target adenosines with minimal bystander editing. **e**, Distribution of secondary structures along the target sequence in top gRNA designs for +3 editing (M671V), 0+1 coediting (K670G), and 0+1+3 co-editing.

We performed HTS utilizing a library of 58,000 gRNA designs with diverse secondary structures based on heuristics derived from natural substrates and ADAR preferences. Although many gRNAs enabled editing the two A-to-I base edits required for the K670G substitution or the single A-to-I base edit required for the M671V substitution, none of the 58,000 designs could edit all three adenosines simultaneously (Fig. 3b).

To identify ADAR substrates that selectively edit all three target adenosines, we trained a CNN that jointly modeled editing efficiency at all three sites (positions 0, +1, and +3) and the specificity score. These models achieved high predictive power, with Spearman’s r > 0.9 for all four metrics. We then usedActMax to design 245 unique gRNAs with high predicted efficiency and specificity at all three sites (Fig. 3c) and validated them experimentally with our HTS. Many designs successfully edited all three adenosines within the same RNA molecule, with 15 gRNAs achieving greater than 40% editing at all three positions (Fig. 3d). These designs combined two crucial structural elements: a 3-4 bp bulge at position +12 observed in 0 and +1 coeditors, and a 1-2 bp mismatch at the target adenosine observed in +3 editors (Fig. 3e).

Although our ML approach successfully discovered novel gRNA designs enabling linked RNA editing for APP, these models remain target specific and are not generalizable across different ADAR substrates (Supplementary Fig. 5). This limitation underscored the need for a target-agnostic model built from sampling across many targets with high sequence and structural diversity.

### A generalizable model for directing ADAR activity

To develop a generalizable model for predicting ADAR activity against any target, we generated a training dataset sampling diverse target and gRNA sequences. We first optimized library design using hairpin T_M_ prediction as a surrogate for secondary structure. Using T_M_ calculation ^38^ as ground truth, we simulated ADAR gRNA hairpin libraries of various size, number of targets, and number of gRNAs per target. We found that approximately 1,000 targets with 10 gRNAs per target could predict the T_M_ of unseen hairpins with a Spearman’s r of 0.82 (Supplementary Fig. 6). Based on this result, we screened a PolyTarget library comprising 50,253 gRNAs across 5,643 targets, with significant nucleotide diversity throughout the entire gRNA-target substrate, including insertions and deletions, and enrichment for mismatches at the target adenosine to promote ADAR editing (Extended Data Fig. 1a) ^38^.

CNN models were trained on a target-stratified subset of 80% of the PolyTarget library and achieved strong predictive power for the 20% of withheld targets across all editing metrics (Figure 4a, Spearman’s r = 0.58 – 0.67). Target primary sequence alone also showed reasonable predictive accuracy (Extended Data Fig. 1b, Spearman’s r = 0.414 – 0.541), reflecting ADAR1 and/or ADAR2’s natural propensity to edit any sequence independent of gRNA design and the larger target-to-gRNA sequence space in the library design. The CNNs further validated their generalization on a separate HTS targeting adenosines at two upstream reading frames of progranulin (GRN uORF1 and GRN uORF2, Spearman’s r = 0.37 – 0.60), indicating we did not overfit the PolyTarget library (Extended Data Fig. 1c).

**Fig 4.**
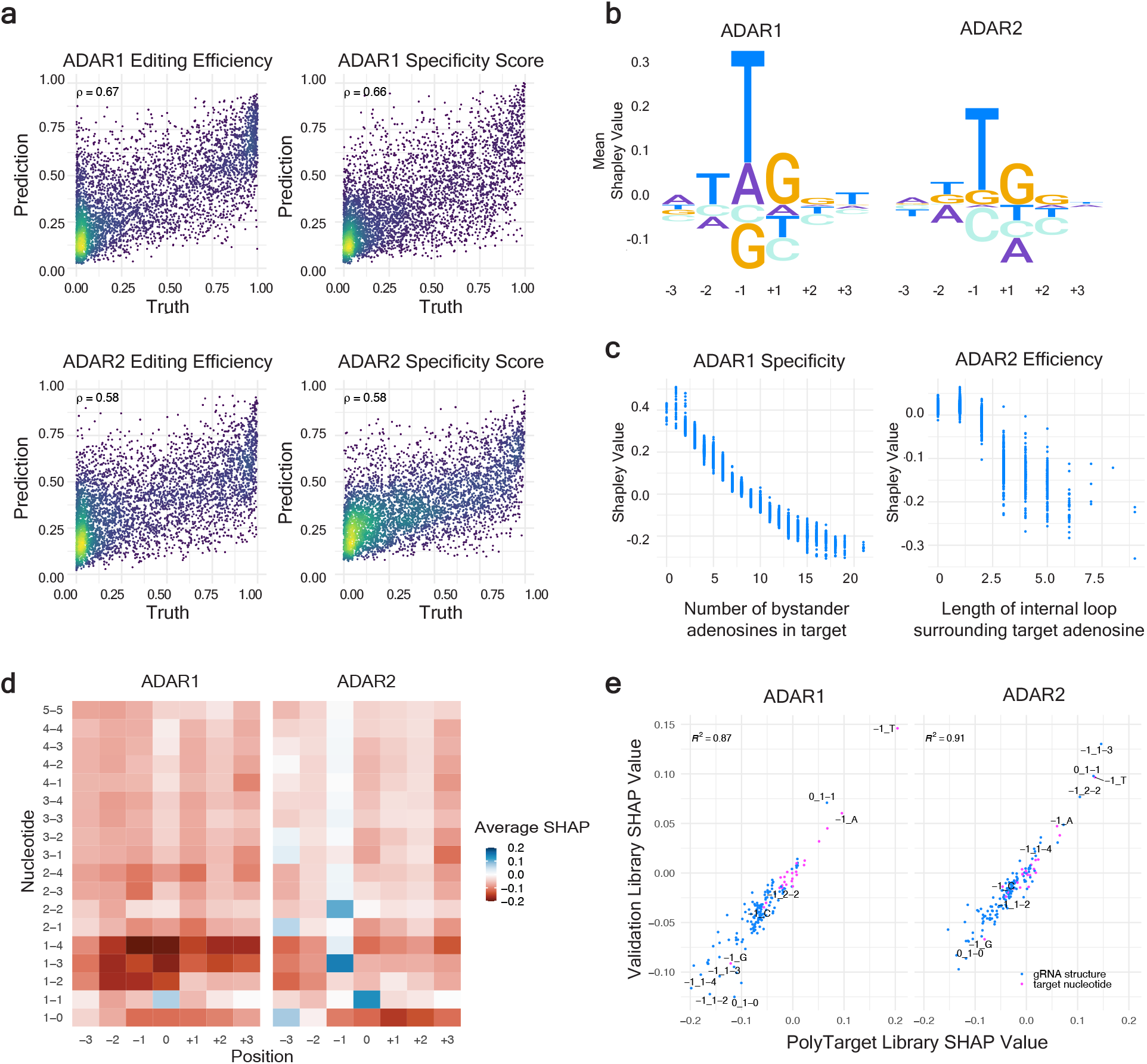
Generalizable model of ADAR activity across substrates. **a**, Predictive performance of CNN ensemble models on held-out targets for editing efficiency and specificity scores for ADAR1 and ADAR2, trained on the PolyTarget library. **b**, SHAP values of three target nucleotides upstream or downstream of each adenosine. **c**, SHAP values showing global characteristics predictive of reduced ADAR editing outcomes. Left: negative correlation between number of bystander adenosines in the target and ADAR1 editing specificity. Right: Correlation between size of internal bulges and loops encompassing on-target adenosine with ADAR2 editing efficiency. **d**, SHAP values of positionally encoded secondary structures representing their predicted marginal contributions to on-target editing across all adenosines in all targets. **e**, Reproducibility of SHAP values estimated on guide structures and target sequences between the PolyTarget library (x-axis) and validation library (y-axis).

To further elucidate the underlying rules governing ADAR editing, we trained an XGBoost model on secondary structure features of the target-guide scaffold and quantified feature importance using SHAP values ^39^. This analysis not only replicated but also expanded upon known ADAR editing preferences, including the 5′-[UA]AG-3′ motif (Fig. 4b) ^10, 11^. Our model revealed distinct preferences between ADAR isoforms, with ADAR1 displaying a more pronounced dispreference for a 5′ G than ADAR2, which could overcome this dispreference when accompanied by a suitable secondary structure. We observed that an increase in the number of bystander adenosines within a target correlated with decreased specificity (Fig. 4c), and internal bulges or loops larger than two nucleotides encompassing the target adenosine were associated with diminished editing efficiency.

Further investigation of isoform-specific preferences (Fig. 4d, e, Extended Data Fig. 2) showed only moderate correlation in on-target adenosine editing between ADAR1 and ADAR2 (*R*^2^ = 0.43). Notable distinctions included ADAR2’s dispreference for a 3′ A, stronger preference for a 2-2 symmetric bulge at the target adenosine, and ability to edit with 1-3 or 1-4 target-guide asymmetric bulges (one target nucleotide unpaired across from three or four gRNA nucleotides). U-deletions 3′ of the target adenosine were associated with a greater reduction in ADAR2 editing compared to ADAR1. We identified optimal secondary structures for promoting ADAR2 on-target editing, including 2-2 symmetric and. 1-3 asymmetric bulges (Fig. 4d). The directionality and magnitude of the effects of these structural characteristics on editing was subsequently validated in a separate library of 37 targets (Fig. 4e).

To evaluate the effect of gRNA structures relative to their distance from the target adenosine, we evaluated sliding windows of 1 to 15 nucleotides around each target adenosine. A three-nucleotide window captured most of the predictive power for on-target editing efficiency compared to the full model incorporating the entire gRNA-target sequence (Supplementary Fig. 7), reinforcing existing knowledge in the field regarding the importance of local sequence context in ADAR editing.

The ability to construct a predictive CNN model from the PolyTarget Library that reflects established ADAR preferences bolstered our confidence in the model’s potential to generate gRNAs de novo for any target in any sequence context.

### DeepREAD: a generative model for gRNA design

To develop generative models for gRNA design, we utilized a CNN paired with ActMax and explored alternative architectures, particularly denoising diffusion probabilistic models (DDPMs). DDPMs have shown promise in image generation and have been successfully adapted for biological applications such as protein and DNA design for regulatory functions ^40, 41^. We trained a bit diffusion model, a DDPM variant suited for discrete RNA sequences, conditioning on target-gRNA sequences and associated editing metrics for ADAR1 and ADAR2 from our PolyTarget library (Fig. 5a). We termed the two modules of ActMax and the bit diffusion-based generative model DeepREAD (Deep Learning for RNA Editing by ADAR Design).

**Fig 5.**
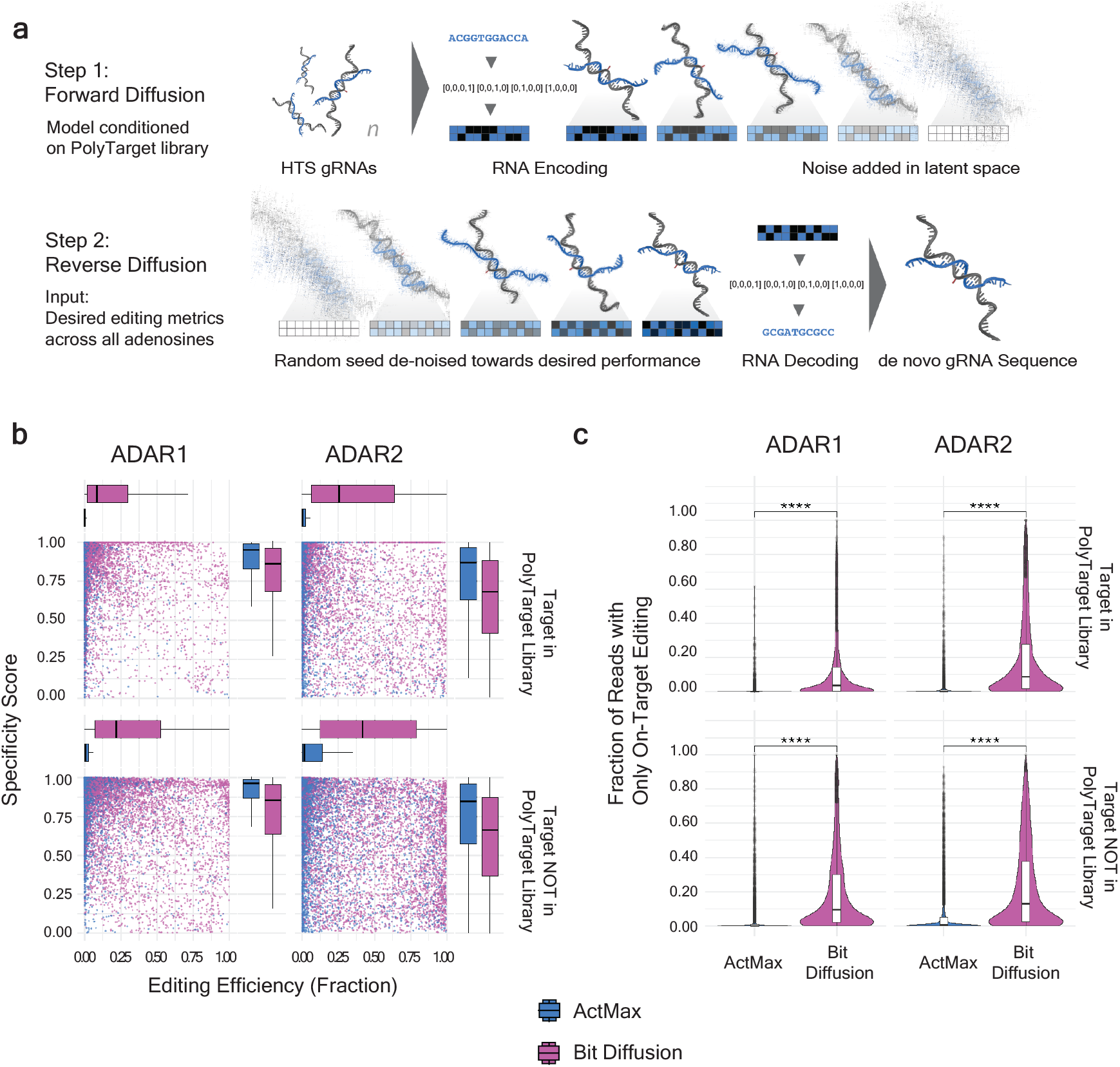
Generative gRNA design for unseen targets. **a**, Schematic of the bit diffusion model used to generate gRNA sequences conditioned to achieve any given editing profile. **b**, Comparison of experimental validation performance of gRNA designs generated by activation maximization (ActMax) and bit diffusion, assessed by on-target editing efficiency and specificity score. Models are both trained on the PolyTarget library data and validated on additional gRNA designs for previously observed targets (top) and unseen targets (bottom). **c**, Distribution of fraction of reads with only a single A-to-G correction at the intended target adenosine for ActMax and bit diffusion designs for previously observed targets (top) and unseen targets (bottom).

To evaluate the efficacy of our modeling approaches, we generated a validation library of gRNAs targeting 37 target adenosines, including 15 from the PolyTarget library and 22 unseen targets, using both bit diffusion and ActMax techniques. We benchmarked these against rational designs, which included an A-C mismatch at the target adenosine or GA-GC bulge at the -1 and target position for adenosines with a 5′ G neighbor, combined with A-G mismatches or U-deletions across from bystander adenosines ^16, 21, 32^.

The bit diffusion model proficiently generated gRNAs where the conditioned input values matched the empirical data for ADAR1 and ADAR2 in the validation library, affirming the model’s accuracy in reflecting each isoforms’ activities (Extended Data Fig. 3a). Additionally, the bit diffusion model generated gRNAs with higher on-target editing but slightly reduced specificity score compared to those derived from ActMax (Fig. 5b). To better capture both on-target editing and specificity in a single metric that would be most reflective of the intended outcome, we quantified sequencing reads with only a single on-target edit and no bystander edits at any other adenosine on the same read (henceforth referred to as fraction on-target only). The fraction on-target only was significantly higher for bit-diffusion-designed gRNAs compared to ActMax, for both the 15 targets that were originally in the PolyTarget library and for the 22 unseen targets (Fig. 5c). The bit diffusion designs also outperformed rational designs and heuristics for improving ADAR editing efficiency and specificity, including A-G mismatches and U-deletions (Extended Data Fig. 3b).

On a target-by-target basis, the bit diffusion model outperformed rational design approaches for generating efficient and specific gRNAs for 35 and 36 of the 37 targets for ADAR1 and ADAR2, respectively (Extended Data Fig. 3c). Of the 37 targets tested, bit diffusion generated gRNAs achieving over 50% on-target only editing for 22 targets with ADAR1 and 30 targets with ADAR2. ActMax surpassed this threshold for far fewer targets (11 for ADAR1 and 22 for ADAR2), while random A-G mismatches or U-deletions performed even worse. For many targets (e.g., KCNB1, HBB, MUTYH, PMM2, and STAT1) bit diffusion was the only strategy to generate efficient designs (Supplementary Fig. 8). These results validated the use of DDPMs, traditionally used for image generation, for effective a priori design of gRNAs that can achieve efficient and specific editing of novel targets.

### DeepREAD generates therapeutically viable gRNAs with species cross-reactivity and allelic specificity

To demonstrate the utility of our DeepREAD model, we generated a panel of gRNAs targeting the MECP2^R168X^ nonsense mutation associated with Rett Syndrome ^42^. A-to-I RNA editing of the opal mutation, UGA-to-UGI, introduces a tryptophan codon that restores translation of the full-length protein ^43^. Rett syndrome is a X–linked dominant disease where roughly half of patient neurons express wildtype MECP2, making it desirable for an RNA editing therapy to correct mutant transcripts but leave wildtype RNA untouched. Additionally, the standard mouse model of this disease carries the human Rett mutation but also two nearby rodent-primate sequence differences. Given these facts, we set two challenging gRNA design criteria for DeepREAD to generate gRNAs with maximum value for therapeutic development: (1) high editing for both mouse and human MECP2^R168X^ transcripts to enable testing across disease-relevant model systems; and (2), reduced editing of human and mouse MECP2^WT^ transcripts to achieve allelic specificity (Fig. 6a). These properties of a gRNA could simplify the clinical path while ensuring functionality and safety across pre-clinical models. We used DeepREAD to generate novel gRNA designs conditioned for high on-target editing and low bystander editing for human MECP2^R168X^. We then used an XGBoost model, also trained on the PolyTarget library, to score these generated gRNAs for predicted on-target editing and specificity of mouse Mecp2^R168X^, human MECP2^WT^ and mouse Mecp2^WT^. Designs were prioritized to maximize predicted specific editing for mouse and human MECP2^R168X^ while minimizing editing of WT transcripts. A panel of 294 designs was selected for testing in an engineered HEK293 cell line expressing all four transcript isoforms of interest. Several gRNAs demonstrated the desired RNA editing activity, with cross-reactive editing of human and mouse MECP2^R168X^ and limited editing of WT transcripts (Fig. 6b), with one gRNA design achieving greater than 51% editing of both mutant transcripts and less than 15% editing of the WT transcripts, while maintaining less than 5% editing for all bystander adenosines (Fig. 6c). This gRNA creates a GA-GC bulge at the target adenosine when hybridized to the human and mouse mutant transcripts, a secondary structure known to facilitate editing of adenosines in a 5′ G context ^16^. Interestingly, due to sequence differences at the - 2 position, this structure is predicted to collapse when the gRNA hybridizes to the WT transcripts, forming an A-bulge that is refractory to ADAR editing ^21^. This provides a unique mechanism for achieving allelic specificity (Fig. 6d).

**Fig 6.**
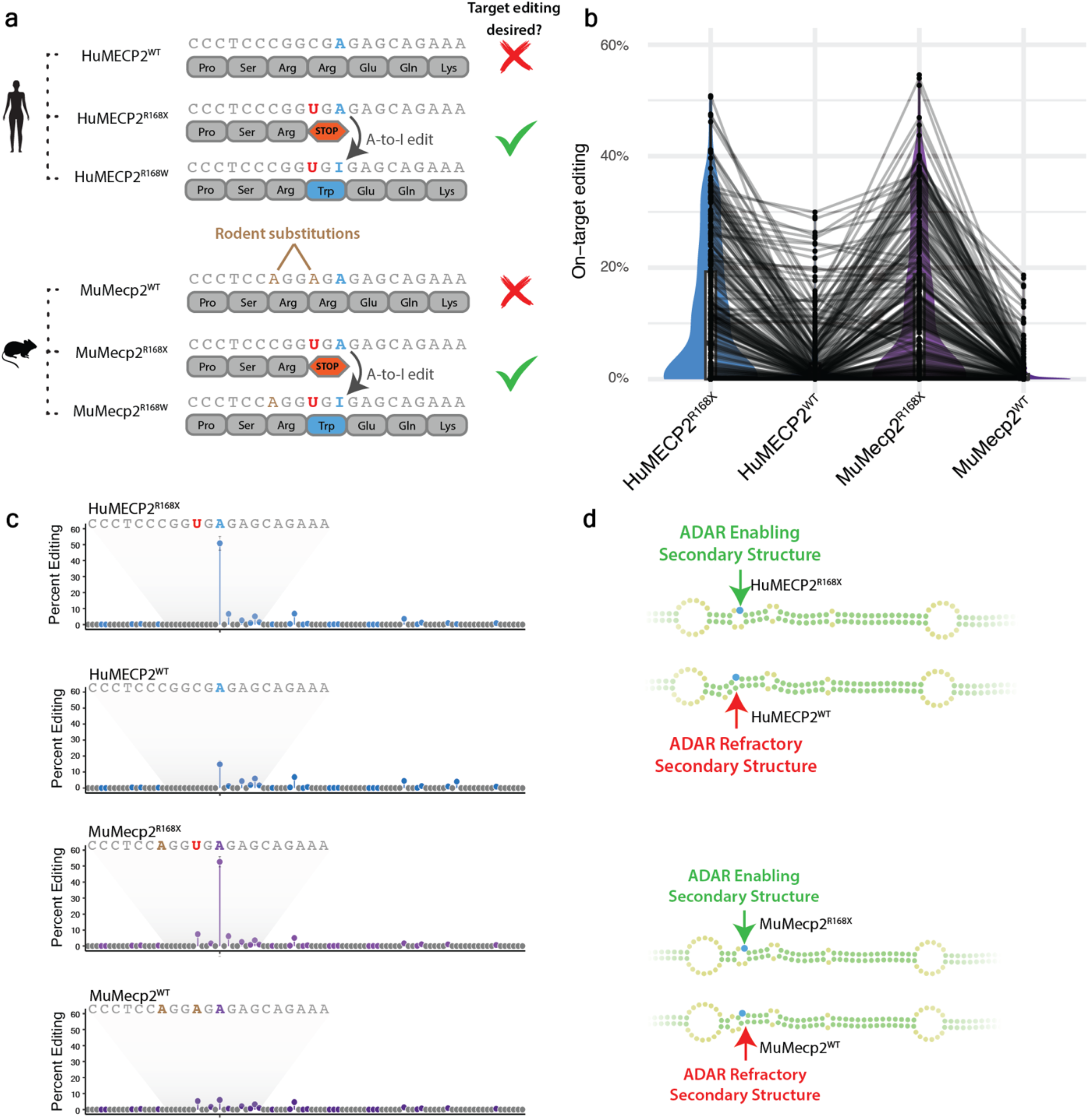
Species cross-reactive and allele-specific gRNAs targeting MECP2^R168X^. **a**, Schematic of the desired gRNA activity profile for optimal therapeutic correction of MECP2^R168X^. **b**, Average percent RNA editing of the target adenosine of the four MECP2 alleles. **c**, Observed HEK293 editing profiles for a generated gRNA, showing percent RNA editing across all adenosines within the gRNA hybridization region for the four MECP2 alleles. **d**, Predicted secondary structure of the top performing gRNA when hybridized to the four MECP2 alleles, with arrows indicating structural alterations that likely allow ADAR to discriminate between alleles.

These outcomes illustrate DeepREAD’s capability to navigate complex gRNA design for intricate targets, potentially accelerating the development of therapeutic gRNAs with favorable clinical attributes. The model’s ability to design gRNAs with both species cross-reactivity and allelic specificity highlights its potential to simplify preclinical testing and enhance the safety profile of RNA editing therapeutics.

## Discussion

In this study, we demonstrate an integrated approach combining HTS and ML to significantly advance gRNA design for ADAR-mediated RNA editing. Our approach overcomes key limitations of current methods and expands the potential applications of RNA editing beyond simple G-to-A mutation correction. By generating diverse libraries with up to 10^6^ gRNA designs per target, we gained unprecedented insight into the complex relationship between RNA structure and ADAR editing efficiency.

Unlike traditional selection methods that require multiple enrichment rounds to identify rare functional solutions, our strategy leverages the fact that most ADAR substrates exhibit some level of editing activity. This allows our HTS to generate rich datasets where nearly every library member provides valuable information. With the varied designs and editing outcomes from these data, we can then learn how to rapidly generate gRNA designs that meet stringent criteria for efficiency and specificity, overcoming ADAR’s natural promiscuity and sequence preferences.

The HTS data enabled training of CNNs that explored vast sequence spaces via ActMax. For the LRRK2 G2019S target, ActMax identified novel gRNA designs with greatly improved editing efficiency and specificity compared to the initial library (Fig. 2). When applied to the more challenging APP target requiring simultaneous editing of three adenosines, ActMax generated designs achieving efficient co-editing where conventional screening failed (Fig. 3). These results demonstrate the power of ML to uncover solutions beyond experimental feasibility.

To develop a generalizable model applicable to any target, we created a PolyTarget library spanning over 5,000 diverse targets. CNNs trained on this dataset showed strong predictive performance on held-out targets (Fig. 4a). These models confirmed known ADAR preferences, such as the 5′-[UA]AG-3′ motif, while also revealing novel insights. For example, we observed that ADAR1 has a more pronounced dispreference for 5′ G compared to ADAR2, which can be partially overcome by suitable secondary structures (Fig. 4c). We also identified optimal structural features for promoting editing across most targets, such as 2-2 symmetric or 1-3 asymmetric bulges encompassing the target adenosine (Fig. 4e).

Although ActMax proved effective for single-target optimization, bit diffusion offered superior performance for de novo gRNA design across unseen targets. DeepREAD, our bit diffusion-based approach, consistently outperformed ActMax and competing rational design strategies in generating efficient and specific gRNAs for previously unseen targets (Fig. 5). A key innovation is DeepREAD’s use of one-hot encoded primary sequences, allowing it to infer structural relationships without sole reliance on free energy minimization algorithms. This enables capture of non-canonical base pairing, which plays a crucial role for ADAR substrates 15.

DeepREAD’s success stems in part from its ability to generate gRNAs with similar MFE ranges as training samples when folded with targets. This leads to more efficient sampling of high-performance design spaces and a higher success rate across diverse targets and ADAR isoforms. DeepREAD inherently learns to integrate complex, varied strategies to minimize bystander editing depending on the sequence context, including G-A mismatches, U-deletions, and G-U wobble base pairs 44. Moreover, DeepREAD can generate gRNAs for any new adenosine in the transcriptome approximately 10,000 times faster than ActMax, allowing rapid production of thousands of gRNAs in minutes for experimental validation. Current efforts to further improve DeepREAD include leveraging transformer architectures to better capture complex RNA structural patterns and dependencies across sequence lengths to focus the model on relevant gRNA elements via attention, using those richer embeddings as inputs into diffusion for generative gRNA design.

To demonstrate the therapeutic potential of our approach, we applied DeepREAD to design gRNAs targeting the MECP2^R168X^ mutation associated with Rett syndrome. We successfully generated designs exhibiting the desired cross-reactivity between human and mouse mutant transcripts while maintaining specificity against WT sequences (Fig. 6). This showcases DeepREAD’s capacity to navigate complex design constraints and potentially simplify preclinical development of RNA editing therapies. Our work significantly expands the potential applications of ADAR-mediated RNA editing beyond simple G-to-A mutation correction. A-to-I editing can induce a wide range of functional changes at both the RNA and protein levels. For instance, targeted editing of regulatory motifs like polyadenylation signals or microRNA binding sites can modulate gene expression ^45, 46^. At the protein level, A-to-I editing can generate 17 different amino acid substitutions, offering unprecedented flexibility in protein engineering. The ability to precisely edit multiple adenosines within a single transcript opens new possibilities for fine-tuning cellular processes. However, further research is needed to fully understand the constraints on multi-site editing within a single gRNA, including the effects of adenosine spacing and periodicity on editing efficiency ^36, 47, 48^. Additionally, the impact of multiple inosine substitutions on translational fidelity and efficiency requires thorough investigation ^44^.

Although our biochemical HTS provides numerous insights into ADAR-mediated RNA editing, it does not capture all factors influencing editing outcomes in cellular contexts, such as the presence of competing RNA binding proteins, target RNA secondary structure, endogenous ADAR levels and modifications, and intracellular pH ^49^. In a cellular context, gRNA expression is also a critical variable, and advancements have been made to leverage novel snRNA expression frameworks or circular gRNAs to drive robust editing ^19, 21, 23^. Encouragingly, DeepREAD-generated gRNAs targeting MECP2 showed high performance in cells despite being trained on these biochemical HTS data. This suggests that ADAR-substrate interactions and the target-gRNA secondary structure are primary determinants of editing outcomes. Future work incorporating high-throughput in-trans gRNA cellular screening and transfer learning approaches will further refine our models for improved in vivo prediction.

In conclusion, our integrated HTS and ML approach represents a significant advance in the rational design of RNA editing therapies. By elucidating complex rules governing ADAR activity and developing powerful generative models like DeepREAD, we have created a platform capable of rapidly generating highly efficient and specific gRNAs for diverse targets. This technology has the potential to accelerate the development of RNA editing-based therapeutics for a wide range of genetic disorders. As the field of RNA editing continues to evolve, we anticipate that the insights and tools presented here will serve as a foundation for further innovation, bringing us closer to realizing the full potential of this promising therapeutic modality.

## Methods

### Protein expression and purification

Recombinant human his-MBP-ADAR2 (UniProt ID: X UP000005640 and His-MBP-ADAR1 (UniProt ID: UP000005640) were expressed and purified by GenScript. ADAR2 was purified using Ni-NTASuperdex 200 16/600 pg His-MBP-ADA, while ADAR1 was purified by a two-step purification (Ni-NTA+ Heparin HP column). ADAR2 was stored in 50 mM Tris-HCl, 500 mM NaCl, 5% Glycerol, pH8 and ADAR1 was stored in 50 mM Tris-HCl, 10% glycerol, 200 mM KCl, 0.5 mM EDTA, 0.01% NP-40, 1 mM DTT for pH8. Purity was > 85% as measured by SDS-PAGE.

### Preparation of the RNA Libraries

For the randomized LRRK2 library, a 177 nt ssDNA library (MW: 54,613; T_M_: 75.6 °C) was synthesized (Genelink) with the underlying sequence “TGCAAAGATTGCTGACTACAGCATTGCTCAGTACTGCTGTAGAAT-Hairpin-ATTCTACAGCAGTACTGAGCAATGCTGTAGTCAGCAATCTTTGCA”. The gRNA region in bold (−8 to +21 from the target adenosine) was spiked at 88% WT and 4% for each alternative base. The library was bottlenecked down to 112k sequences using digital droplet PCR (ddPCR) with PrimeSTAR GXL DNA Polymerase (TAKARA #R050A) to enhance sequencing coverage of designs. All other RNA libraries were synthesized with unique barcode identifiers by Twist Biosciences. The ssDNA oligo libraries were amplified with the PrimeSTAR GXL DNA Polymerase with a T7 Promoter for in vitro transcription (IVT), which was performed using the HiScribe T7 Quick High Yield RNA Synthesis kit (NEB #E2050S) at 37 °C for 16 h. RNA was DNase treated and purified using the Monarch RNA Cleanup Kit (NEB #T2040S).

### ADAR Reactions

The RNA libraries were denatured starting at 95 °C, then cooled by 5 °C increments every 30 s until reaching 37 °C. The final reaction buffer contained 17 mM Tris H-Cl pH8.0, 60 mM KCl, 16 mM NaCl, 2 mM EDTA, 0.003 (v/v) NP-40, 1 mM MgCl2, 1 mg/mL yeast tRNA and 0.5 mM DTT, and RNAseOUT (2 U/mL). Recombinant ADAR was added to a final concentration of 100 nM, and reactions were incubated at 30 °C or 37 °C for 30 min or 60 min, unless otherwise noted. The ADAR reactions were heat-inactivated at 95 °C for 5 min before cooling to 4 °C.

### Library Preparation for Sequencing

After the ADAR reaction, the RNA libraries were reverse transcribed using AMV Reverse Transcriptase (NEB #M0227L) with a primer targeting a universal priming site present on all sequences in the RNA library. The cDNA underwent two rounds of PCR amplification using PrimeSTAR GXL DNA Polymerase to introduce Illumina adapters for sequencing (Illumina #20027213). The libraries were sequenced on an Illumina HiSeq or NextSeq 1000.

### Methods for Validating LRRK2 and MECP2 gRNAs in HEK293T cells

The HEK293T-LRRK2^G2019S^ cell line was engineered using the PiggyBac transposon system to express a minigene containing a portion of the LRRK2^G2019S^ transcript. The HEK293T-MECP2^4iso^ cell was engineered using the PiggyBac transposon system to express the full-length MECP2 open-reading frame with a C-terminal Flag-tag for four variants: human MECP2^R168X^, human MECP2^WT^, mouse Mecp2^R168X^, and mouse Mecp2^WT^. gRNAs were cloned into a proprietary vector under the control of a mouse U7 snRNA promoter and 3′ regulatory elements. Engineered HEK293T cells were seeded 24 h before transfection in 96-well plates. Transfections were performed using TransIT-293 Transfection Reagent (Mirus #2704) according to the manufacturer’s instructions. Cells were harvested 48 h post-transfection, and RNA was isolated using the RNeasy Mini Kit (Qiagen #74104). Reverse transcription was carried out with gene-specific primers and/or oligo dT utilizing SuperScript™ IV Reverse Transcriptase (Thermo Fisher #18090050). Amplicons were amplified utilizing KAPA HiFi DNA Polymerase (HotStart and Ready-Mix formulation) (Roche #07958935001). All samples for LRRK2 and MECP2 were prepared for NGS with Illumina Nextera Unique primers and sequenced on the Illumina iSeq.

### Data analysis

Sequencing read quality was assessed using fastp (version 0.22.0). Reads failing quality thresholds were filtered out, and the remaining reads analyzed using a custom regular expression to quantify editing at each adenosine. Editing levels were calculated as the number of reads with a guanosine at a given position divided by the total number of reads covering that position. On-target editing was estimated at the intended target adenosine, and a specificity score was calculated by the following formula: ((fraction on-target reads + 1) / (sum of bystander reads + 1)). Empirical Bayes shrinkage was applied using the eBayes R package to adjust estimates for targets with low sequencing coverage by borrowing information within each experimental condition (e.g., ADAR1).

### RNA secondary structure prediction and featurization

The secondary structure formed by the hybridization of each gRNA and target sequence was predicted using the Vienna RNAfold package (version 2.5.0a5). RNAfold uses a dynamic programming algorithm to find the secondary structure with the minimum free energy (MFE) for each input sequence ^38^. The predicted structures were represented in dot-bracket notation, where paired bases are denoted by matching parentheses and unpaired bases by dots. Custom scripts were then used to extract relevant features from the predicted secondary structures, such as the number and size of loops, bulges, and stems, as well as the positions of specific structural elements relative to the target adenosine.

### Supervised machine learning

Gradient boosted decision trees were trained using the XGBoost library in Python (version 3.8) and R (version 4.0.3). The input data was split into training and validation sets using an 80/20 split, stratified by target to ensure balanced representation. Hyperparameters were optimized using Bayesian optimization with three-fold cross-validation within the training set. To interpret the trained models, feature importance was estimated using Shapley Additive Explanations (SHAP) ^39^ with the TreeExplainer method, representing the average marginal contribution of each feature to the model’s predictions across all possible feature orderings.

### Generative gRNA design by activation maximization

Sequences representing the self-annealing RNA structure formed by gRNA-target hybridization were one-hot encoded and used as input to train an ensemble of 20 convolutional neural networks (CNNs). The vector representation used as input to the model concatenated the target and gRNA sequence, enabling the evaluation of a gRNA’s performance in relation to its hybridization target. The target sequences all had a length of 41 ribonucleotides, while the gRNA sequences varied in length from 32 to 49 ribonucleotides.

The CNNs were trained to predict editing efficiency, specificity, and MFE, with architectures and hyperparameters selected by random search to minimize validation loss. The hyperparameter search space included the number of layers (4-8), number of channels per layer, kernel size, stride, padding, and dilation. No residual connections were used, and each CNN had a unique configuration. The model endpoints were editing efficiency and specificity (experimentally obtained) and MFE (calculated using ViennaRNA)^38^. The loss function was the mean squared error between the predicted and target values.

To generate novel designs, the trained ensemble performed ActMax ^50, 51^. The network parameters and target portion of the input were frozen, and stochastic gradient descent was used to optimize the gRNA sequence. The optimization objective was defined to minimize a loss function combining the absolute error between the predicted and target values for on-target editing efficiency and specificity (equally weighted), with MFE assigned a weight of zero. The optimization was run for 1,000 iterations with two constraints: (1) the optimized gRNA sequence values were clipped to the range [0,1] and required to sum to one; and (2) the iterative gRNA solution was projected back to the feasible space every 25 steps. The final solution for a given initialization was the nearest one-hot encoding of a sequence after a predetermined number of steps.

To mitigate the generation of gRNAs that highly deviate from the duplex structure when the underlying predictive model exhibited insufficient performance, a constraint was introduced to penalize such deviations. Incorporating a term into the cost function necessitated the addition of a scalar value to balance the two terms. Through repeated evaluation and adjustment of the search space, a parameter value of 0.03 was found to facilitate ActMax in providing gRNAs that balanced the competing aims of desired editing and specificity while minimizing deviations from the duplex.

For the multi-target library, 100 “positive” gRNAs and 10 “negative” gRNAs were selected from pools of designs generated by ActMax and bit diffusion scored by XGBoost models on the conditions of ADAR1, ADAR2, or ADAR1 & ADAR2, based on the product of their editing and specificity predictions. This combined score encouraged the judicious addition of features that enhance specificity while minimally reducing editing efficiency.

### Generative gRNA design by conditional diffusion models

Conditional diffusion models, renowned for their capacity to generate near-realistic images and other modalities, serve as potent tools for selectively sampling from known distributions ^52^. In the realm of gRNA generation, we employ a method akin to one-hot encoding, where each sequence entry assumes a value between -1 and 1 ^41, 53^. Operating in a continuous space, the model accepts discrete values as input and thresholds its outputs back into the discrete domain. The conditional diffusion model operates within the gRNA space, with the conditioning signal containing the one-hot encoded target sequence, and scalar values describing the editing and specificity.

During training, the forward diffusion process iteratively applies Gaussian noise to the sequence’s one-hot encoding over a predetermined number of time steps. In the reverse diffusion process, a denoising U-net model then refines these noised gRNAs, aiming to match the final iteration’s output with the initial input. This process enables the model to learn the distribution defining the interaction between the gRNA and target, as well as the experimentally measured editing and specificity. In each iteration, the gRNA is refined to align with the underlying distribution that governs its complementarity with the target sequence, along with the desired editing and specificity levels. When performing inference, the model samples Gaussian noise and takes as input the set of desired conditioning values before applying the previously described reverse diffusion process to obtain a sample. This novel application of diffusion models to gRNA generation provides a method to efficiently sample from regions of interest (e.g., gRNAs with high editing and specificity).

The dataset was partitioned into 60% training, 20% validation, and 20% test subsets. To prevent data leakage, the splits were stratified based on the targets, ensuring that the model needed to find solutions for unseen targets. The CNN was trained using a stochastic gradient optimizer, while the conditional diffusion model employed an Adam optimizer. The latter was trained for 100 epochs with a learning rate of 1e-4. To stabilize the model and mitigate epoch-to-epoch fluctuations, an exponential moving average was applied with a beta of 0.995. Additionally, the conditional diffusion model utilized a linear beta learning rate scheduler with a beta value starting at 0.0001 and ending at 0.2.

## Acknowledgements

We thank the following individuals for their support and contributions to this work: Katie Rupp and Jason Dean for developing the initial bioinformatics pipeline for the high-throughput screening analysis; Joanne Boysen, Lauren Swanson, and Aditya Radhakrishnan for their assistance with in-cell editing pre-processing analysis; Aydin Abiar for engineering the bit diffusion pipeline; Henry Lee for generating in cell data for LRRK2 gRNAs; Shivani Patel and Anupama Lakshmanan for generating the HEK293 cell line engineered to express all four MECP2 transcript isoforms of interest; Andrew Sadowski and Scott Rich for generating in cell data for MECP2 gRNAs; Susan Byrne and Stephen Burleigh for developing the gRNA expression system; and Stephen Burleigh for scientific discussion during the conceptualization of the biochemical screening system.

## Author contributions

Y.J. and L.R.B. contributed equally to this work as the senior authors. B.B. and A.W.B. conceptualized and developed the HTS. B.B. and L.G. developed the bioinformatics pipeline for analyzing the HTS data. L.R.B. further developed the HTS platform and generated all HTS data. L.G., Y.J., and B.S.B. conceptualized and developed the methods for target-conditioned DeepREAD using activation maximization. B.S.B., Y.J., and R.J.H. developed the methods for target-agnostic DeepREAD using bit diffusion.

Y.C. and Y.J. generated machine learning designs targeting MECP2^R168X^. Y.A.S., B.B., and L.R.B. contributed to the design of gRNA architecture for grafting HTS substrates into full-length gRNAs for expression in cells. B.B. and R.J.H. wrote the manuscript and shared in the intellectual supervision of the work. All authors reviewed and edited the manuscript.

