## Supplementary Figures for "Generative machine learning of ADAR substrates for precise and efficient RNA editing"

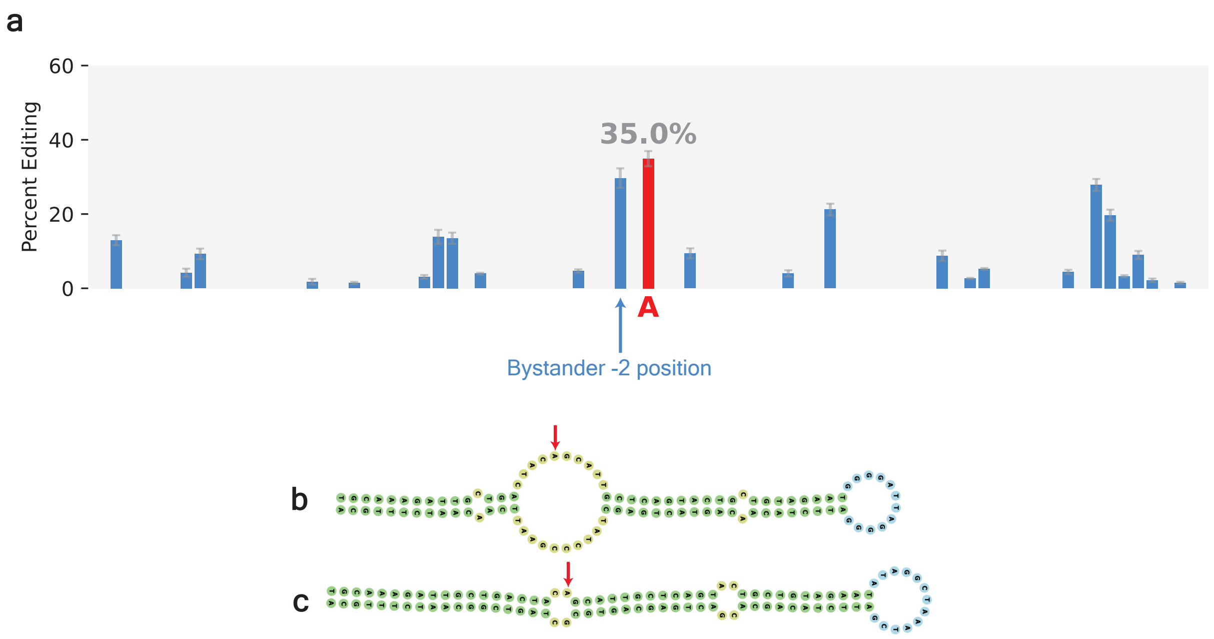


**Supplementary Fig. 1** | Editing profiles and secondary structures of LRRK2 G2019S gRNA designs. **a**, Editing profile of the canonical A-C mismatch gRNA design of LRRK2 G2019S, showing substantial bystander editing at the -2 position. b, Predicted secondary structure of a highly specific and efficient LRRK2 G2019S substrate identified in the biochemical HTS. c, Predicted secondary structure of a ML-derived LRRK2 G2019S with an unconventional “G” placed opposite the target adenosine. When grafted into a full-length gRNA, this design exhibited on-target editing only in the presence of exogenous ADAR2 expression.

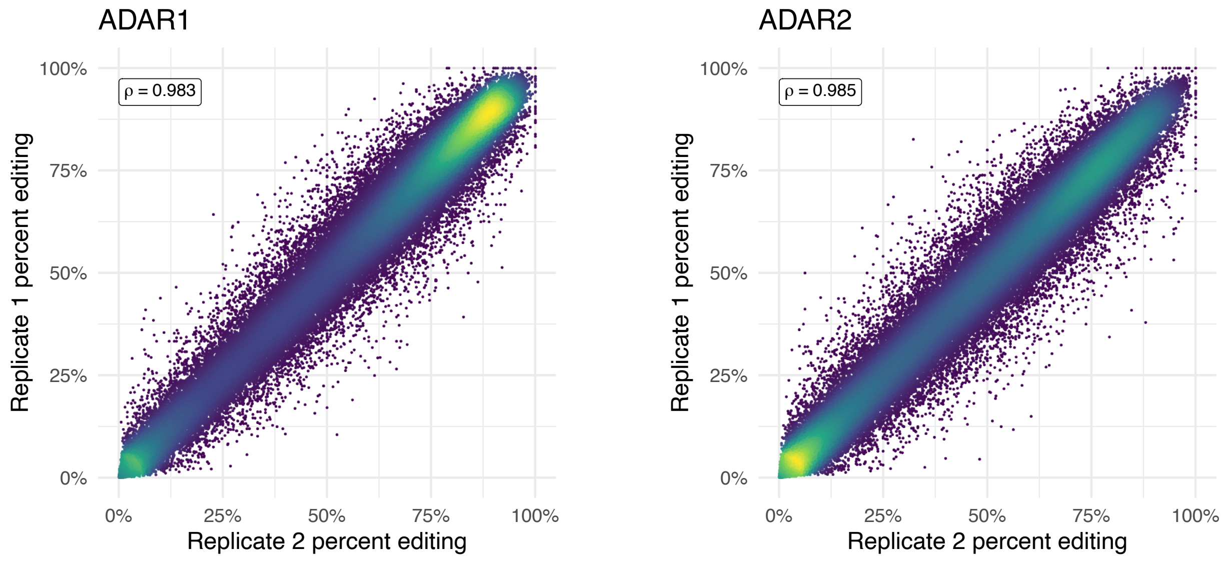


**Supplementary Fig. 2** | Technical reproducibility of the biochemical HTS assay on the LRRK2 G2019S library.

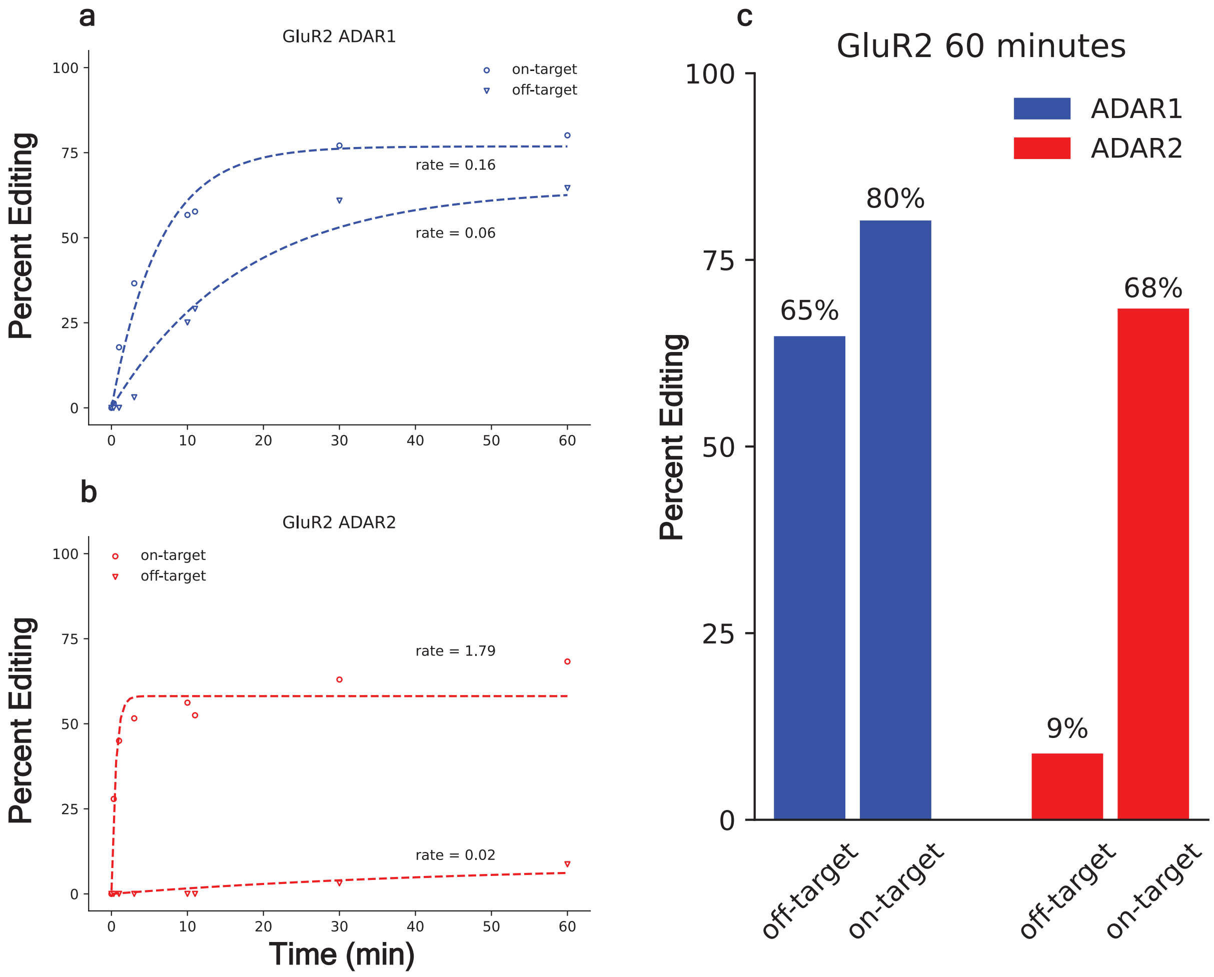


**Supplementary Fig. 3** | Deamination kinetics of the GluR2 R/G hairpin substrate. **a**, **b**, Curve fits of editing fraction time course analysis using four-parameter logistic regression for ADAR1 (**a**) and ADAR2 (**b**) respectively. c, Editing fraction of on-target and the 5′ neighboring adenosine after 60 minutes.

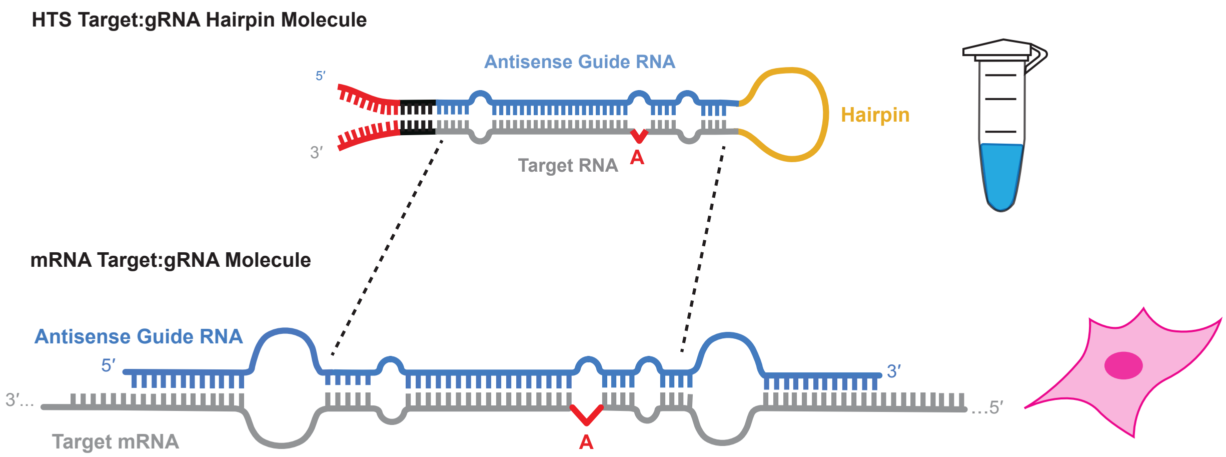


**Supplementary Fig. 4** | Schematic representation for the in-cell validation process.

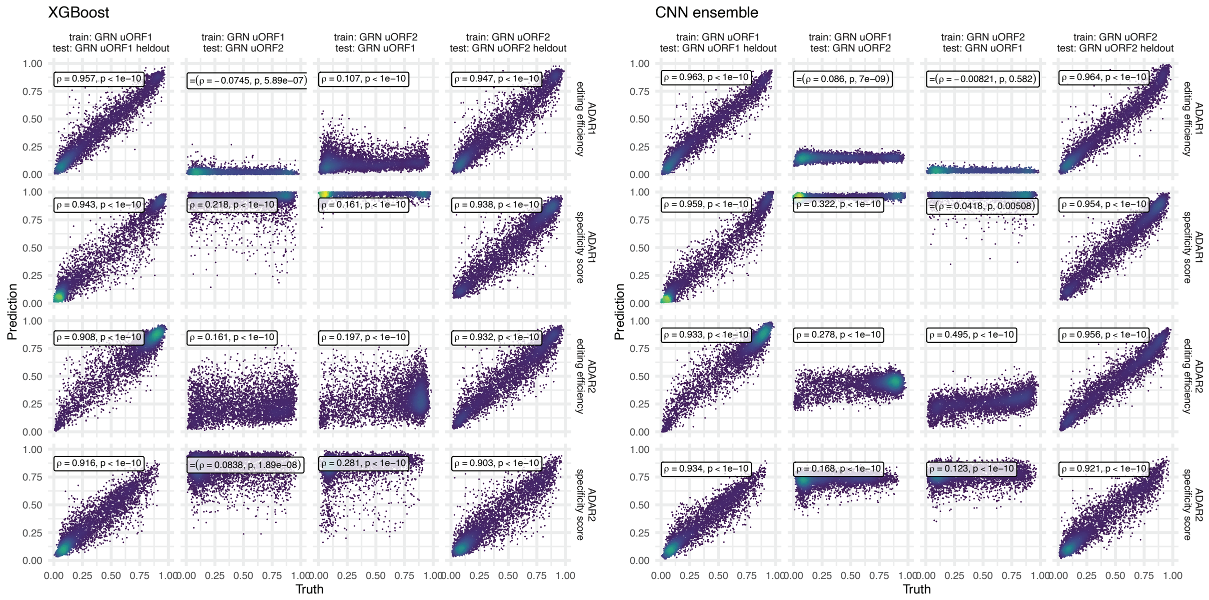


**Supplementary Fig. 5** | Limited generalizability of models trained on single-target datasets. Performance of XGBoost and CNN ensemble models trained on individual target HTS datasets (GRN uORF1 or GRN uORF2) and tested on held-out data from the same or the other target.

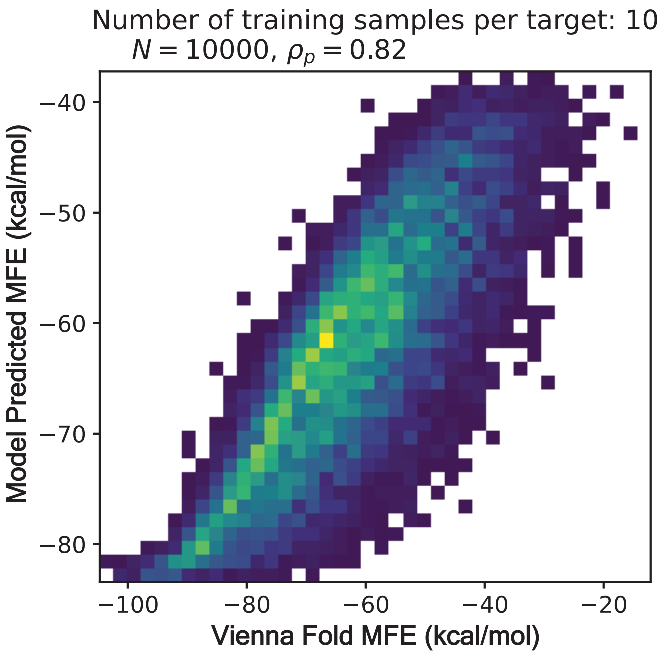


**Supplementary Fig. 6** | XGBoost prediction of minimum free energy (MFE) versus ground truth (Vienna Fold predictions), using synthetic dataset of 10,000 targets with 10 gRNAs per target.

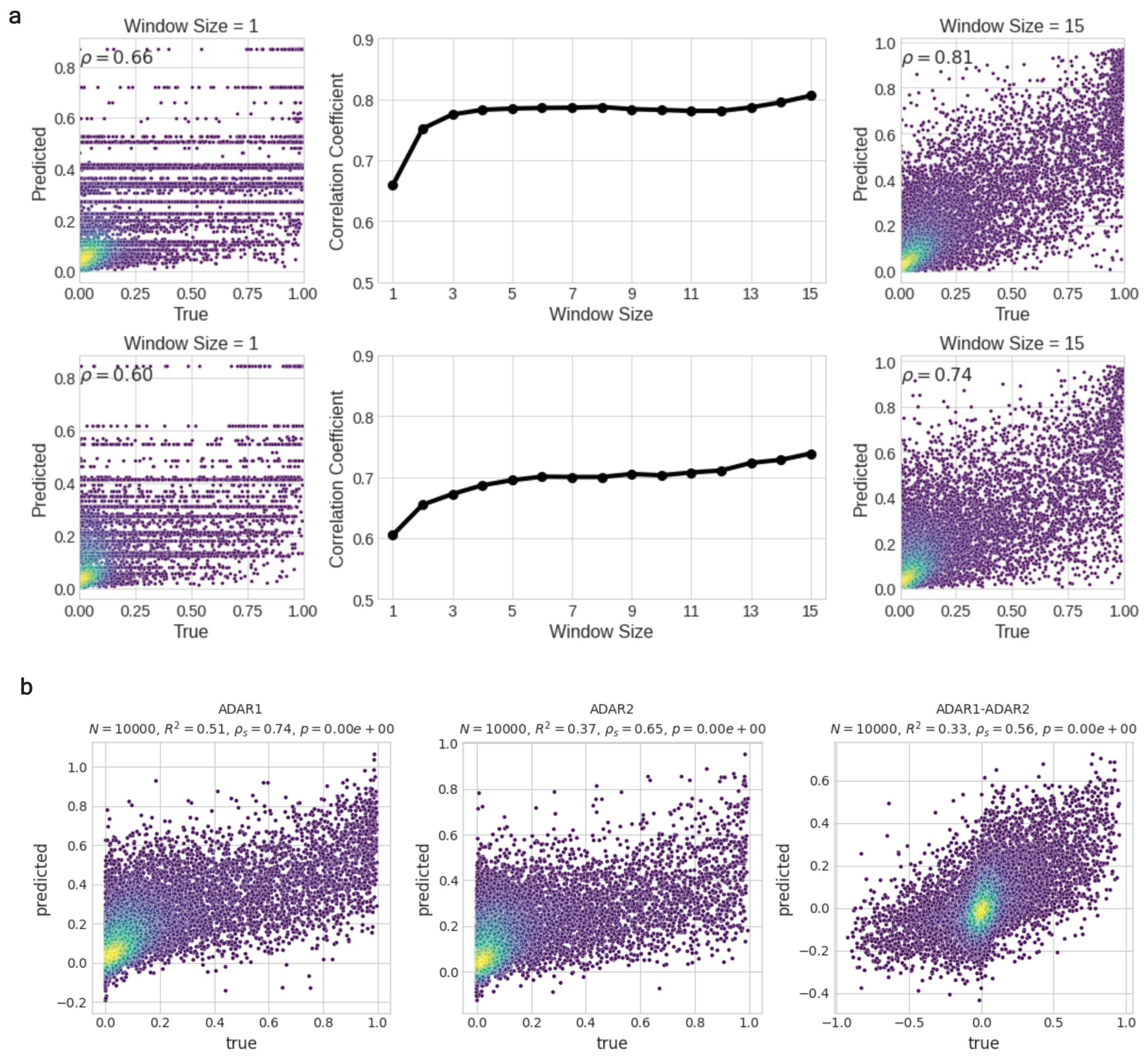


**Supplementary Fig. 7** | Evaluating the sequence information as a function of nucleotides surrounding relative to each adenosine. **a,** Performance of XGBoost models as a function of the number of nucleotides surrounding each adenosine in the target:guide duplex for ADAR1 (top) and ADAR2 (bottom). **b,** Predictive performance of XGBoost models using a 7-nucleotide window around each adenosine along the target:guide duplex.

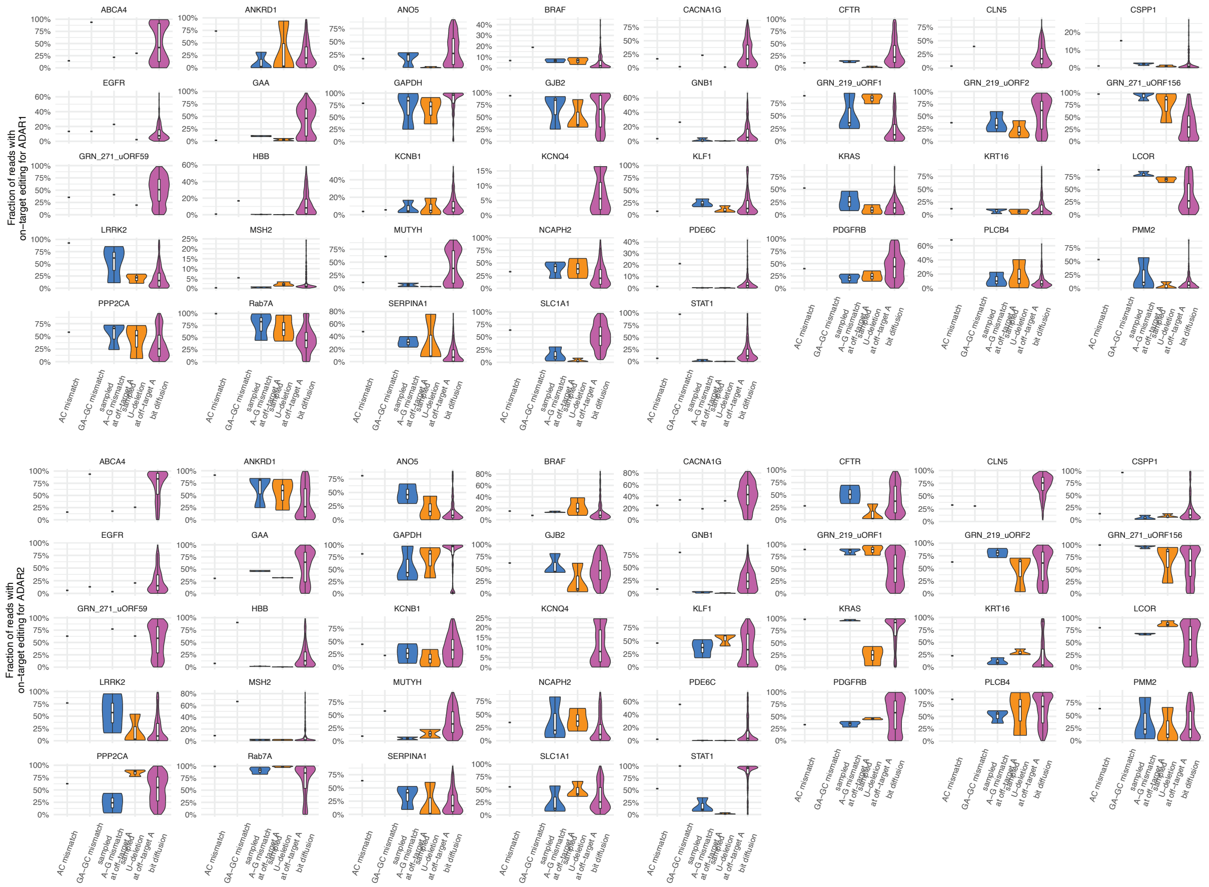


**Supplementary Fig. 8** | Distributions of edited read fractions at on-target adenosines. Comparison of editing outcomes across 37 targets for guides generated using different strategies: A-C mismatch, GA-GC mismatch, sampled A-G mismatches at bystander adenosines, sampled U-deletions at bystander adenosines, or bit diffusion. Results are shown for both ADAR1 (top) and ADAR2 (bottom).
