## Extended Data for "Generative machine learning of ADAR substrates for precise and efficient RNA editing"

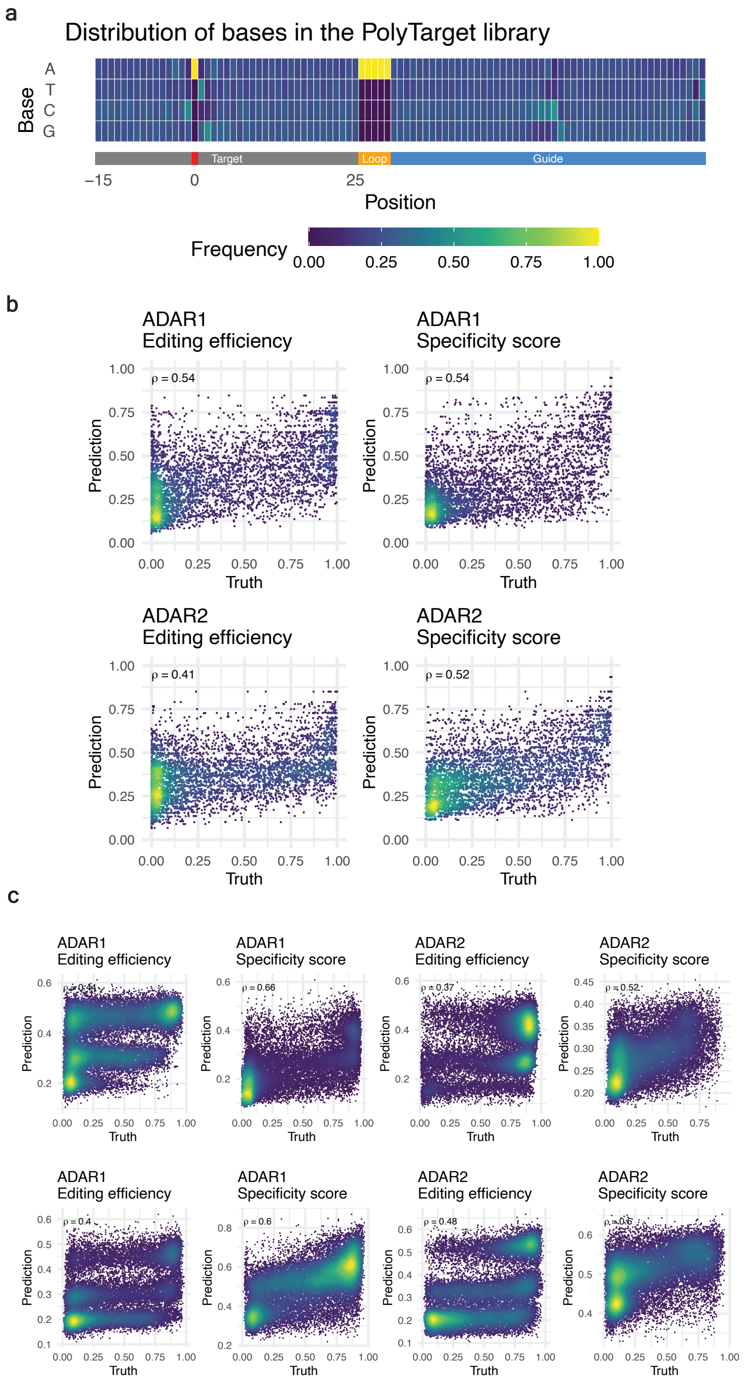


**Extended Data Figure 1 | Characterization of the PolyTarget library. a,** Nucleotide diversity across the library. b, Correlation between predicted and observed editing outcomes based on a CNN ensemble trained only on target sequences. c, Correlation between predicted and observed editing outcomes from a CNN ensemble trained on the PolyTarget library for two unseen targets, GRN uORF1 and GRN uORF2.

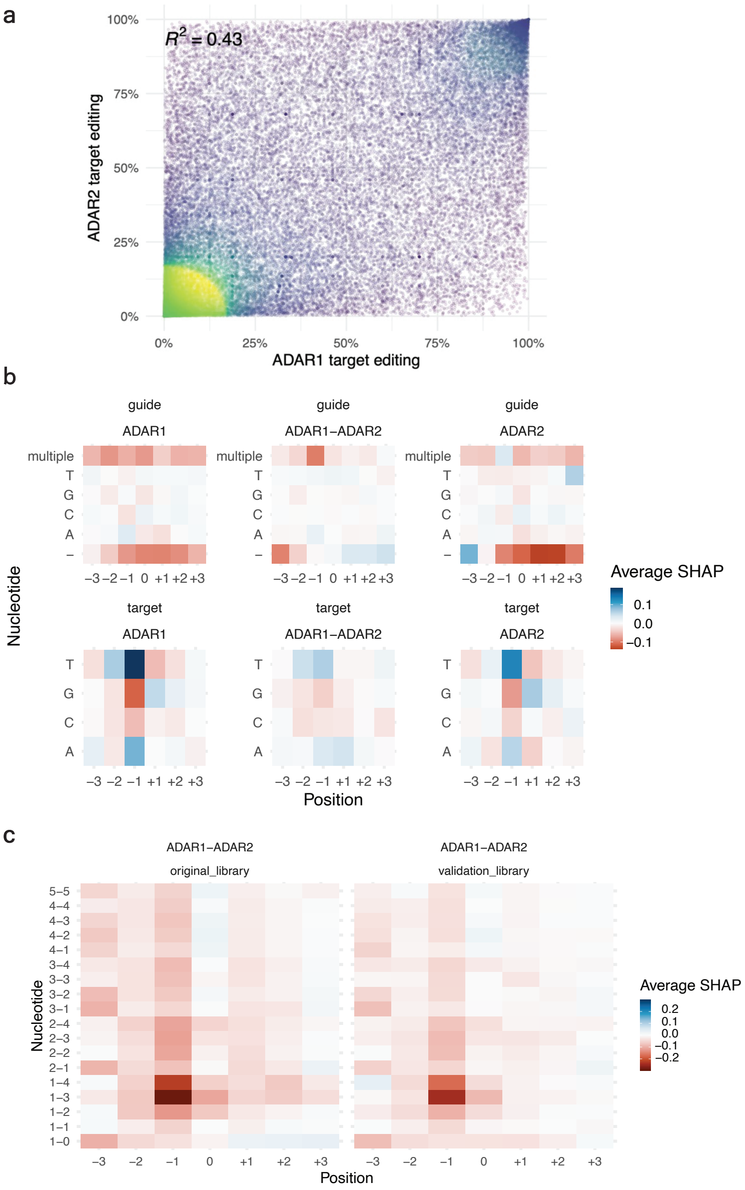


**Extended Data Figure 2 | ADAR isoform differences in editing efficiency.** a, Observed correlation between on-target adenosine editing outcomes for ADAR1 and ADAR2 from the PolyTarget library. b, SHAP values estimated from three nucleotide adenosine windows around each adenosine for either the guide (top) or target (bottom) sequence on editing efficiency. "multiple" indicates more than one gRNA nucleotide across from a target nucleotide position, and "-" indicates a deletion across from the corresponding target position. "ADAR1-ADAR2" refers to the difference between editing efficiency for ADAR1 and ADAR2 for each adenosine. **c.** SHAP values of positionally encoded secondary structures representing their predicted marginal contributions to on-target editing across all adenosines in all targets for the difference between ADAR1 and ADAR2 editing efficiency in the original PolyTarget library and subsequent validation library.

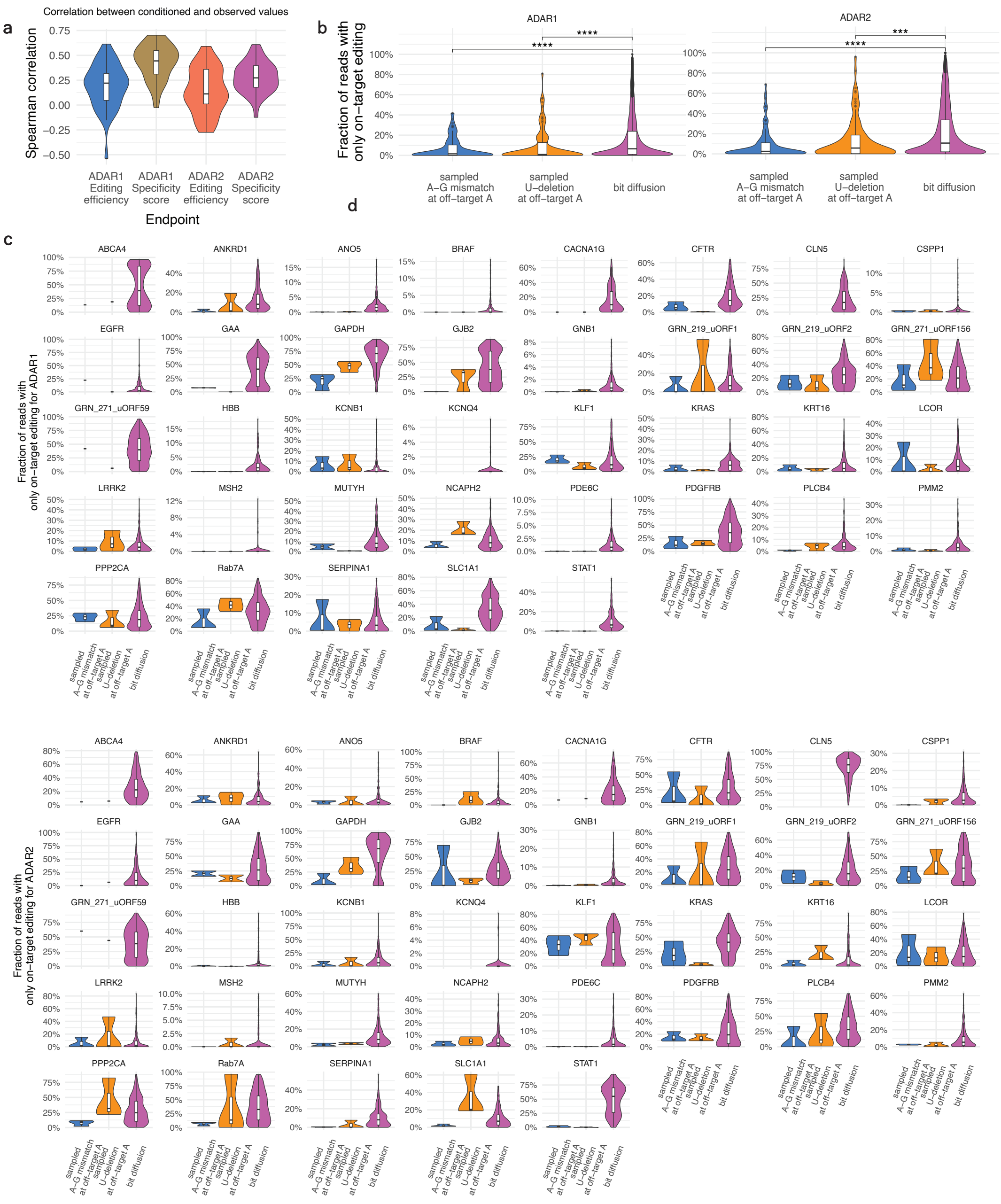


**Extended Data Figure 3 | Comparison between gRNA design strategies for unseen targets.** a, Fraction of reads with only on-target editing for gRNA generated by bit diffusion, random sampling of A-G mismatches, or random sampling of U-deletions across bystander adenosines. b, Correlation between conditioning values used to generate bit diffusion gRNAs and observed editing performances. c, Distributions of fractions of reads with only a single A-to-G correction at the intended target adenosine across 37 targets from guides generated with either an A-C mismatch, GA-GC mismatch, sampled A-G mismatches at bystander adenosines, sampled U-deletions at bystander adenosines, or through bit diffusion for both ADAR1 (top) and ADAR2 (bottom).
